# Partitioning The Effects of Eco-Evolutionary Feedbacks on Community Stability

**DOI:** 10.1101/104505

**Authors:** Swati Patel, Michael H Cortez, Sebastian J Schreiber

## Abstract

A fundamental challenge in ecology continues to be identifying mechanisms that stabilize community dynamics. By altering the interactions within a community, eco-evolutionary feedbacks may play a role in community stability. Indeed, recent empirical and theoretical studies demonstrate that these feedbacks can stabilize or destabilize communities, and moreover, that this sometimes depends on the relative rate of ecological to evolutionary processes. So far, theory on how eco-evolutionary feedbacks impact stability exists for only for a few special cases. In our work, we develop a general theory for determining the effects of eco-evolutionary feedbacks on stability in communities with an arbitrary number of interacting species and evolving traits for when evolution is slow and fast. We characterize how eco-evolutionary feedbacks lead to stable communities that would otherwise be unstable, and vice versa. We show how this characterization provides a partitioning of the roles of direct and indirect feedbacks between ecological and evolutionary processes on stability, and how this partitioning depends on the rate of evolution relative to the ecological time scales. Applying our methods to models of competing species and food chains, we demonstrate how the functional form of trade offs, genetic correlations between traits, and the rate of evolution determine whether eco-evolutionary feedbacks stabilize or destabilize communities.

## INTRODUCTION

In recent years, there has been growing empirical evidence highlighting that eco-evolutionary feedbacks can dramatically impact community-level properties, including stability (Pimentel et al. 1963; Yoshida et al. 2003; Becks et al. 2010; Sanchez and Gore 2013; Steiner and Masse 2013; Kasada et al. 2014). Moreover, there is evidence that evolution happens at various rates, ranging from slower than to commensurate to ecological rates (Darimont et al. 2009; terHorst 2010; DeLong et al. 2016; Hendry 2016), and this affects the impact of feedbacks (Becks et al. 2010; Turcotte et al. 2011; Reznick 2013). Theoretical work to mechanistically understand how eco-evolutionary feedbacks impact stability has mostly focused on specific ecological modules with just one or two evolving species (Abrams and Matsuda 1997; Fussmann et al. 2007; Cortez and Ellner 2010; Vasseur et al. 2011; Hendry 2013; Cortez 2016). Here, we present a general framework to determine the role of eco-evolutionary feedbacks on stability, accounting for the relative time scales of ecological and evolutionary processes.

Determining when an ecosystem of interacting species is stable is of fundamental importance in ecology, offering insight into how the ecosystem will respond to inevitable perturbations. A stable system is able to withstand small but frequent perturbations (Schreiber 2006). Understanding how eco-evolutionary feedbacks impact stability is even more important given the rise of anthropogenic changes, including climate change and habitat disturbances, which fundamentally perturb ecosystems. These perturbations are likely to lead not only to ecological responses, i.e., population density changes, but also drive an evolutionary response, i.e., trait changes in the populations, due to changing selection pressures (Darimont et al. 2009; Hendry et al. 2011; Lankau et al. 2011; Alberti 2015). Indeed, in a meta-analysis, Darimont et al. (2009) showed that trait change was more rapid in human-disturbed ecosystems than from natural causes. As ecosystems are perturbed, evolutionary trait changes in the enclosed populations and subsequent feedbacks with the ecological dynamics can play a critical role in determining how ecosystems respond to perturbations.

Previous work has explored how eco-evolutionary feedbacks affect community stability but this work has been limited to either small, specific communities with one or two evolving species or numerical rather than analytical results. In particular, analyses of predator-prey (Abrams and Matsuda 1997; Cortez and Ellner 2010; Cortez 2016), two competing species (Vasseur et al. 2011), three-species apparent competition (Schreiber et al. 2011; Schreiber and Patel 2015), and three-species intraguild predation (Patel and Schreiber 2015) models highlight that the stabilities of eco-evolutionary coupled systems depend on the relative time scales of the ecological and evolutionary process. Theoretical studies such as these are important given the empirical evidence that evolution can happen at varying time scales relative to ecological processes (DeLong et al. 2016). In more complex communities, the increased number of direct and indirect interactions make understanding the impacts of eco-evolutionary feedbacks on stability more challenging (Hendry 2016) and thus far, studies have predominantly used numerical techniques (Kondoh 2003; Barabás and D’Andrea 2016), which can sometimes limit their ability to provide a mechanistic understanding. Hence, a general analytical theory of when and how eco-evolutionary feedbacks alter stability in general ecological communities is lacking.

In this paper, we address this challenge through a new approach for determining stability in models that couple ecological and evolutionary dynamics. In particular, we present analytical conditions to infer stability for slow and fast evolution in a general model for coupled eco-evolutionary dynamics and describe the role eco-evolutionary feedbacks have on these conditions. Specifically, we are able to partition the effects of ecology (direct effects of density changes on population dynamics), evolution (direct effects of trait changes on selection) and eco-evolutionary feedbacks (indirect effects of density changes on population dynamics mediated by trait changes and indirect effects of trait changes on selection mediated by population density changes) on stability, enabling a more mechanistic understanding of stability in communities with ecological and evolutionary dynamics. Importantly, we show that these feedbacks can fundamentally change predictions on the qualitative dynamics compared to predictions when ecology and evolution are uncoupled. To demonstrate the utility of our approach, we apply it to models of two competing species with one species evolving in one trait and a food chain with one species evolving in multiple traits.

## MODEL

To evaluate the effects of eco-evolutionary feedbacks on stability, we examine a general multi-species model that couples population dynamics with evolutionary trait dynamics. We consider *k* species interacting, with population densities *N*_1_ *… N_k_*, and *ℓ* total ecologically-important evolving traits with trait values as *x*_1_ *… x_ℓ_*. The model is

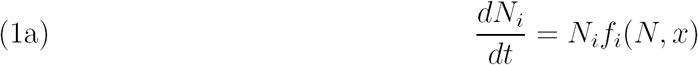

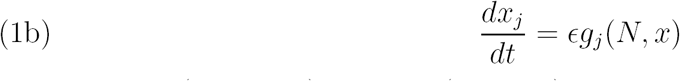

where *N* = (*N*_1_ *… N_k_*) and *x* = (*x*_1_ *… x_ℓ_*). *f*_*i*_ represents the per-capita fitness of species *i* and describes how the growth of species *i* depends on the population densities of all species and all traits. *g*_*j*_ represents the selection function and describes how trait *j* evolves due to the selective pressures imposed by all population densities and traits.

Finally, *ϵ* determines the time scale separation between the ecological and evolutionary dynamics. When *ϵ* is small, the evolutionary process occurs very slowly relative to the ecological processes (hereafter referred to as “slow evolution”). For example, slow evolution occurs when there are little fitness differences among individuals and low levels of genetic variance, which constrain evolutionary potential. When *ϵ* is large, the evolutionary process occurs very quickly relative to the ecological processes (hereafter referred to as “fast evolution”). For example, fast evolution, relative to ecology, occurs when species densely occupy an environment, which constrains their growth. Hence, population densities do not change much over many generations, while traits may change. Indeed, many classical population genetics models of evolution and coevolution assume an extreme version of this scenario in which population densities remain constant while genotypic frequencies change (Lande 1976; Seger 1988; Gavrilets 1997; Nuismer and Doebeli 2004). The limit of fast evolution also serves as a useful heuristic tool for understanding eco-evolutionary dynamics when the time scale separation is less extreme or absent (Cortez and Ellner 2010; Cortez and Weitz 2014; Cortez 2015). Notably, this model form has the flexibility to incorporate any number and combination of species interactions, including predator-prey, mutualistic, and competitive interactions. In particular, we make no initial assumptions about the functional form of growth rates *f*_*i*_, which determine the dynamical consequences of the species interactions. This model form also has the flexibility to incorporate any number of evolving traits, including coevolution between multiple species and multitrait evolution within one species. Our modeling of the evolutionary dynamics implicitly assumes:

(1) for each trait, evolution can be represented by changes in a single continuous quantity (e.g. the mean of a quantitative trait or the frequency of an allele) all changes in these continuous quantities are attributed to evolutionary selection

These assumptions are met by a number of eco-evolutionary modeling frameworks, e.g., adaptive dynamics (Marrow et al. 1996; Geritz et al. 1998) and quantitative genetic approaches based on Lande (1976). To demonstrate the flexibility of our framework, we describe two particular applications of the Lande approach. For example, when one quantitative trait is evolving in each species, with constant genetic variances, and selection is frequency independent, the evolutionary dynamics are given by

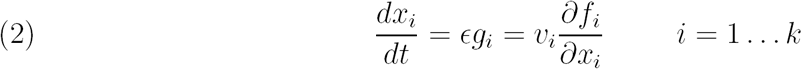

where *x*_*i*_ and *v*_*i*_ are the mean and genetic variances of the evolving trait, respectively. Note that the trait changes in the direction of increasing fitness due to the selection gradient. When all the genetic variances and the selection gradient are sufficiently small relative to growth when measured on the same scale, evolution is slow and this generates a natural time scale separation between the ecological and evolutionary processes.

Alternatively, when one species, say species *i*, is evolving in multiple quantitative traits, the evolutionary dynamics are given by

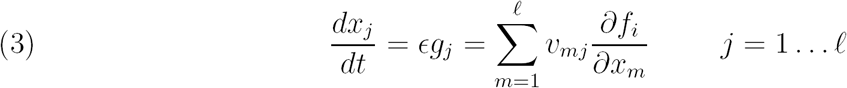

where *x*_*j*_ is the mean of the *j*-th evolving trait and *v*_*mj*_ is the genetic covariance between trait *m* and *j*. Here, selection acting on one trait affects the evolutionary dynamics of any other covarying trait and hence, may affect the eco-evolutionary dynamics (Lande and Arnold 1983; Kopp and Matuszewski 2013).

## EQUILIBRIUM STABILITY AND ECO-EVO FEEDBACKS

How a system at equilibrium responds to perturbations determines its stability. In general, an equilibrium is stable if the system returns to the equilibrium following small density or trait perturbations. Equilibrium stability is determined by the eigenvalues of the Jacobian matrix. In particular, for differential equation models, if the stability modulus, i.e. the largest real part of the eigenvalues, of the Jacobian is (positive) negative then the equilibrium is (un)stable. For any matrix, *M*, we use *s*(*M*) to denote the stability modulus.

The Jacobian for (1) has the form

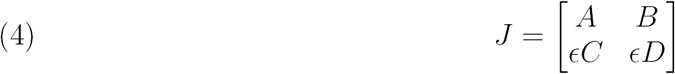

where *A, B, C,* and *D* are submatrices and *ϵ* is as in (1). These submatrices represent different components of the eco-evolutionary dynamics of the community. Submatrix *A* captures the direct effects ecological processes have on species densities (Figure 1 I); it has elements 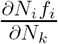, i.e. how the growth rate of species *i* depends on the density of species *k*. Commonly referred to as the community interaction matrix (Levins 1968; May 1973; Pimm and Lawton 1978; Bender et al. 1984; Yodzis 1988), it is the Jacobian of the ecological dynamics and determines stability in the absence of evolutionary dynamics. We call the system *ecologically stable* if *s*(*A*) *<* 0; when traits are fixed, population densities return to equilibrium following a perturbation. Analogously, *D* captures the direct effects of evolutionary processes on trait dynamics and has elements 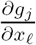, i.e. how selection on trait *j* depends on trait *ℓ*. We call the system *evolutionarily stable* if *s*(*D*) *<* 0; when population densities are fixed, traits return to equilibrium following a perturbation. Submatrix *B* captures the effects of evolution on ecology; it has elements 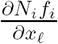, i.e. how the growth rate of species *i* depends on trait *ℓ*. *C* captures the effects of ecology on evolution; it has elements 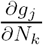, i.e. how selection on trait *j* depends on the density of species *k*.

**FIGURE 1.**
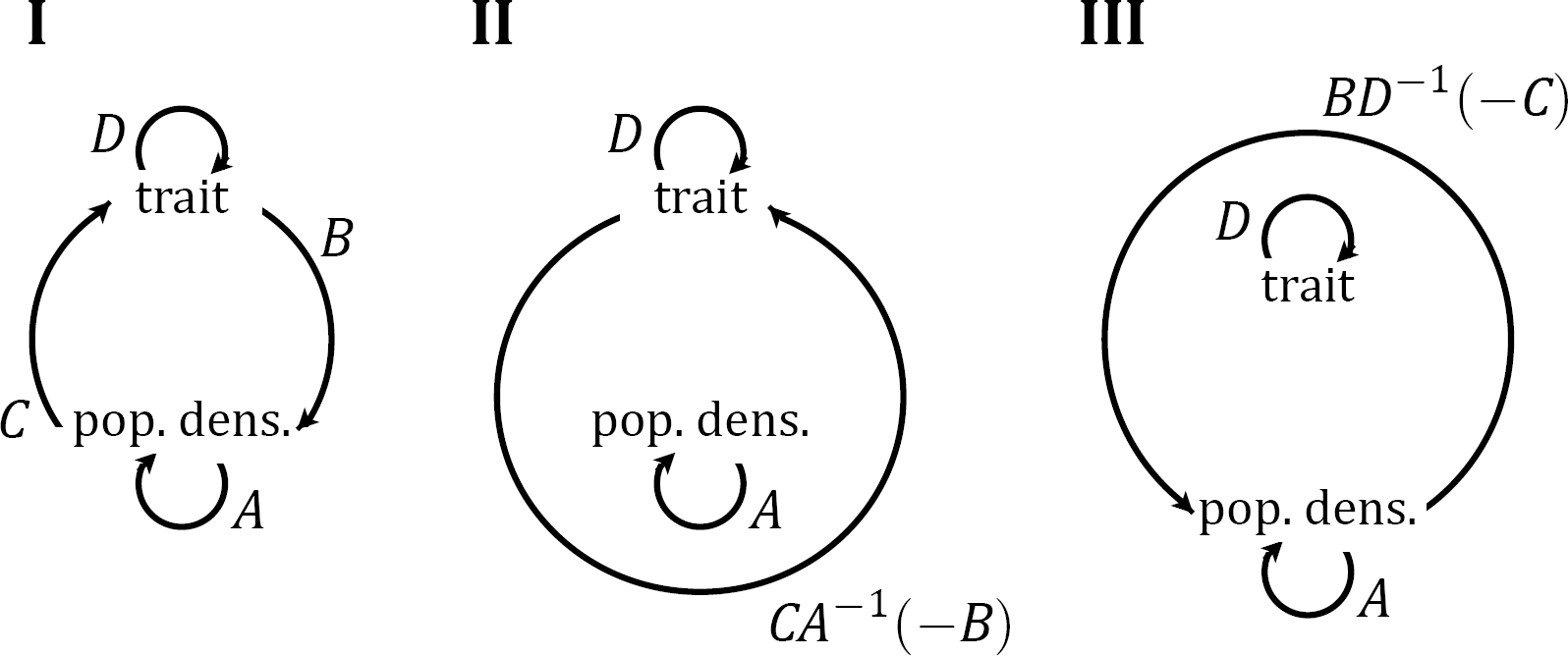
Schematics showing important feedbacks in coupled eco-evolutionary systems. Traits and population densities have direct effects (clockwise arrows) on themselves and on each other (I). These generate the indirect evo-eco-evo feedback (counterclockwise arrow in II) and the indirect eco-evo-eco feedback (counterclockwise arrow in III). If there is a time scale separation between the ecological and evolutionary dynamics, (II) and (III) show the critical feedbacks for determining stability when evolution is slow and fast, respectively.

The equilibrium of model (1) is *overall stable* if *s*(*J*) *<* 0 and *overall unstable* if *s*(*J*) *>* 0. When ecological and evolutionary dynamics are uncoupled, *B* and *C* are zero matrices (implying no effects of ecology on evolution and vice versa) and only the direct effects captured in *A* and *D* determine stability. In this case, *s*(*J*) is the maximum of *s*(*A*) and *s*(*D*). Hence, the stability of the whole system is determined by the stabilities of the separate ecological and evolutionary components. An equilibrium must be both ecologically stable and evolutionarily stable to be overall stable (i.e. *s*(*J*) *<* 0 if and only if *s*(*A*) *<* 0 and *s*(*D*) *<* 0).

On the other hand, when ecological and evolutionary dynamics are coupled, indirect eco-evolutionary feedbacks between population densities and traits also play a role in overall stability. Coupling of ecological and evolutionary processes yields two indirect feedback loops that are important for overall stability when evolution is slow or fast: (1) the evo-eco-evo feedback and (2) the eco-evo-eco feedback. In the evo-eco-evo feedback, changes in traits cause changes in population densities, which in turn alter selection on the traits (Figure 1 II). Mathematically, the three steps are captured by *CA*^*−1*^(*−B*). In the first step of the evo-eco-evo feedback, a change in the traits drives changes in population growth, given by *B*. In the second step, the population densities respond to this change in growth and reach a new equilibrium, given by *A*^*−1*^(*−B*). This expression was used by Levins (1968), Bender et al. (1984), and Yodzis (1988) to describe how population densities in communities respond to external perturbations (Δ*N*). In the final step, changes in population densities drive changes in selection pressures, given by *C*, closing the feedback loop. Similarly, in the eco-evo-eco feedback, changes in population densities cause changes in the traits, which in turn alter population dynamics (Figure 1 III). The corresponding three steps are captured by the matrix *DB*^*−1*^(*−C*).

## GENERAL STABILITY RESULTS FOR SLOW AND FAST EVOLUTION

When and how do these direct and indirect feedbacks affect the stability of equilibria when ecology and evolution are coupled? To address this question, we discuss the stability conditions for cases when there is a time scale separation between ecological and evolutionary processes. The time scale separation enables us to derive analytical conditions that partition the effects of ecology, evolution and eco-evolutionary feedbacks on stability. Mainly, we show that the evo-eco-evo feedback (Figure 1 II) is critical for stability when evolution is slow and that the eco-evo-eco feedback (Figure 1 III) is critical for stability when evolution is fast. We begin with the case of a single slowly-evolving trait coupled to multi-species population dynamics to present and describe the intuition behind the results.

### Communities with a single slowly evolving trait

We consider a community with *k* interacting species, in which one focal species is evolving in a single trait. Throughout this section, we assume that the evolutionary process is slow relative to the ecological processes (*ϵ* is small). In this case, the Jacobian for an equilibrium point can be represented by

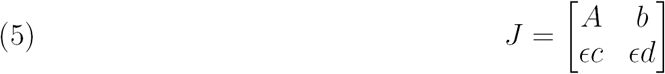

where *b* and *c* are column and row vectors, respectively, with length equal to the number of species and *d* is a scalar. Then, the eco-evolutionary equilibrium is stable if the equilibrium is ecologically stable (i.e. *s*(*A*) *<* 0) and

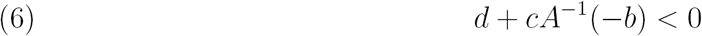

In words, stability requires that the sum of the direct evolutionary feedback and the indirect evo-eco-evo feedback is negative. Hence, if the feedback is negative (*cA*^*−1*^(*−b*)), then the more negative it is, the more it has a stabilizing effect. Alternatively, if the feedback is positive, then the more positive it is, the more it has a destabilizing effect. A proof of this condition is given in Appendix A1.

Intuitively, the stability conditions describe a response to perturbation that proceeds in two steps: a fast ecological response, which leads to the condition *s*(*A*) *<* 0, and a slow evolutionary response, which leads to the condition (6). In the fast response, the population densities either quickly converge to a population equilibrium (ecological stability) or diverge from it (ecological instability). Hence, stability in the fast (ecological) dynamic is always necessary.

In the slow response, the trait slowly changes, due to the direct evolutionary effects. These slow trait changes drive the population densities to continuously respond and quickly approach a new equilibrium corresponding to the changed trait. This in turn affects the selection pressures, giving rise to the indirect evo-eco-evo feedback. When condition (6) is met, these direct and indirect effects act in conjunction to drive the traits and population densities to return to the eco-evolutionary equilibrium (i.e., overall stability; Figure 2A).

**FIGURE 2.**
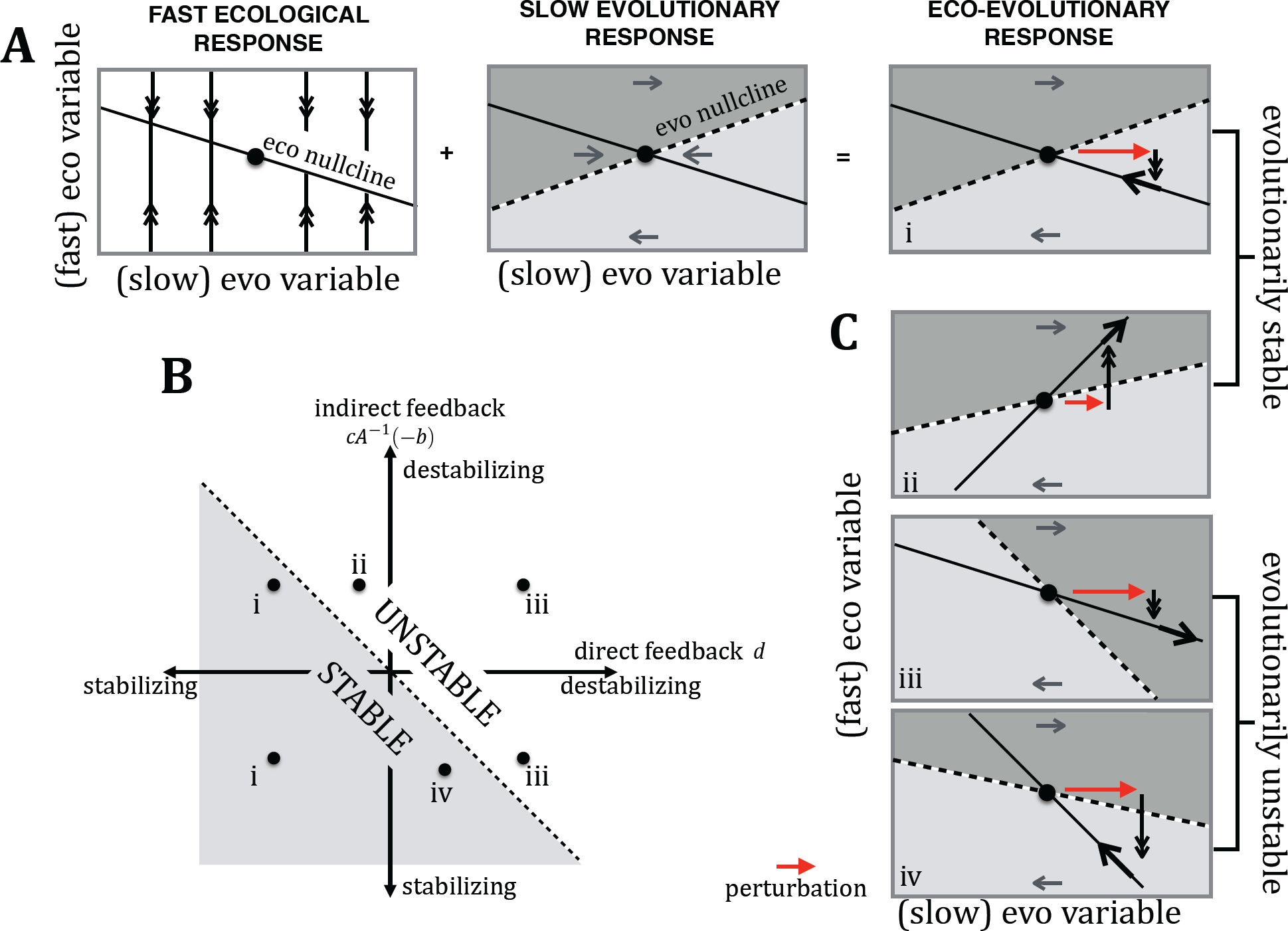
Visual representation of the stability results for a community with a single slowly evolving trait. In (A), the three schematic plots show the fast ecological dynamics (double black arrows in first plot), the slow evolutionary dynamics (gray arrows in second plot), and their joint response to a perturbation (red arrow in third plot). In (B), the dotted line depicts when the sum of the direct (*d*) and the indirect evo-eco-evo feedback effect (*cA*^*−1*^(*−b*)) is zero, which separates stable from unstable. The points correspond to the four possible scenarios discussed in the text: evolutionarily stable and overall stable (i), evolutionarily stable and overall unstable (ii), evolutionarily unstable and overall unstable (iii), and evolutionarily unstable but overall stable (iv). The two-step eco-evolutionary responses for each of these scenarios are shown in the schematic plots in (A) and (C).

From this condition, an equilibrium can be characterized by one of four possible cases, differentiated by whether it is evolutionarily stable as well as whether it is overall stable in the coupled system. Specifically, an equilibrium can be (i) evolutionarily stable and overall stable, (ii) evolutionarily stable and overall unstable, (iii) evolutionarily unstable and overall unstable, and (iv) evolutionarily unstable and overall stable (see Figure 2). Interestingly, scenario (ii) highlights that an equilibrium that is ecologically stable and evolutionarily stable need not be overall stable; this occurs when the indirect evo-eco-evo feedback is sufficiently destabilizing. Conversely, scenario (iv) highlights that even if an equilibrium is evolutionarily unstable, it can be stabilized when the evo-eco-evo feedback is sufficiently stabilizing to counter the destabilizing effect of the direct evolutionary feedbacks. In this latter case, we say that the evo-eco-evo feedback stabilizes the equilibrium.

### Communities with multiple slowly evolving traits

Suppose now we wish to model a community, in which there are multiple evolving traits. This is appropriate for when many species are each evolving in a single trait (as in equation 2), a single species is evolving in multiple traits (as in equation 3) or, most generally, when many species are evolving in multiple traits. When there are multiple evolving traits, collectively in all the species, the response to perturbations and the conditions for equilibrium stability are nearly identical to the single trait case. In particular, the eco-evolutionary equilibrium is stable if the equilibrium is ecologically stable (i.e., *s*(*A*) *<* 0) and

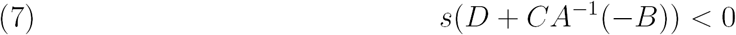

For details, see Appendix A1. Analogous to the previous section, the fast ecological response leads to condition *s*(*A*) *<* 0 and the slow evolutionary response leads to the condition (7) for stability. The key difference between the multiple trait and single trait cases is that when there are multiple traits, the direct evolutionary feedback (*D*) and the indirect evo-eco-evo feedback are represented by matrices (*CA*^*−1*^(*−B*)). Hence, stability is determined by the stability modulus of this matrix sum. As with a single evolving trait, an equilibrium that is ecologically stable and evolutionarily stable can be unstable due to the evo-eco-evo feedbacks. On the other hand, evo-eco-evo feedbacks can stabilize an equilibrium, even if it is evolutionarily unstable.

### Communities with multiple fast evolving traits

To understand the effects of eco-evolutionary feedbacks when evolution is fast, we study the opposite limit (large *ϵ*). For example, this may occur when population densities are constrained near a carrying capacity and their growth is limited. The eco-evolutionary equilibrium is stable if the equilibrium is evolutionarily stable (i.e., *s*(*D*) *<* 0) and

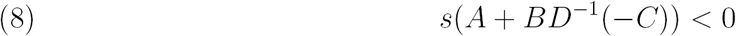

Equation (8) shows that when evolution is fast, equilibrium stability requires that the direct ecological effects (*A*) in addition to the indirect eco-evo-eco feedbacks (*BD*^*−1*^(*−C*)) are negative. Note that when the ecological variables are only one-dimensional, this condition simplifies to *a* + *bD*^*−1*^(*−c*) *<* 0 (analogous to the case of a single slowly evolving trait).

When evolutionary processes are fast relative to ecological processes, the two-step response to perturbations is reversed: the evolutionary response is fast, which leads to the condition *s*(*D*) *<* 0 and the ecological response is slow, which leads to condition (8). In the fast response, provided the equilibrium is evolutionarily stable, traits quickly approach the trait equilibria, due to the direct evolutionary effects. In the slow response, the population densities slowly change, due to the direct ecological effects. These density changes drive the traits to continue to change, which in turn affects the growth of each population, giving rise to the indirect eco-evo-eco feedback. When condition (8) is met, these direct and indirect effects act in conjunction to drive the traits and population densities to return to the eco-evolutionary equilibrium.

## APPLICATIONS TO TWO COMPETING SPECIES

Here, we apply our stability conditions to determine how eco-evolutionary feedbacks can alter stability between two competing species. Classic two-species competition theory asserts that a coexistence equilibrium is ecologically stable if each species competes more strongly intraspecifically than interspecifically (Tilman 2007). Conversely, if both species competes more strongly interspecifically than intraspecifically, then the equilibrium is unstable and the two species do not coexist; initial densities determine which species is competitively excluded (Figure 3).

**FIGURE 3.**
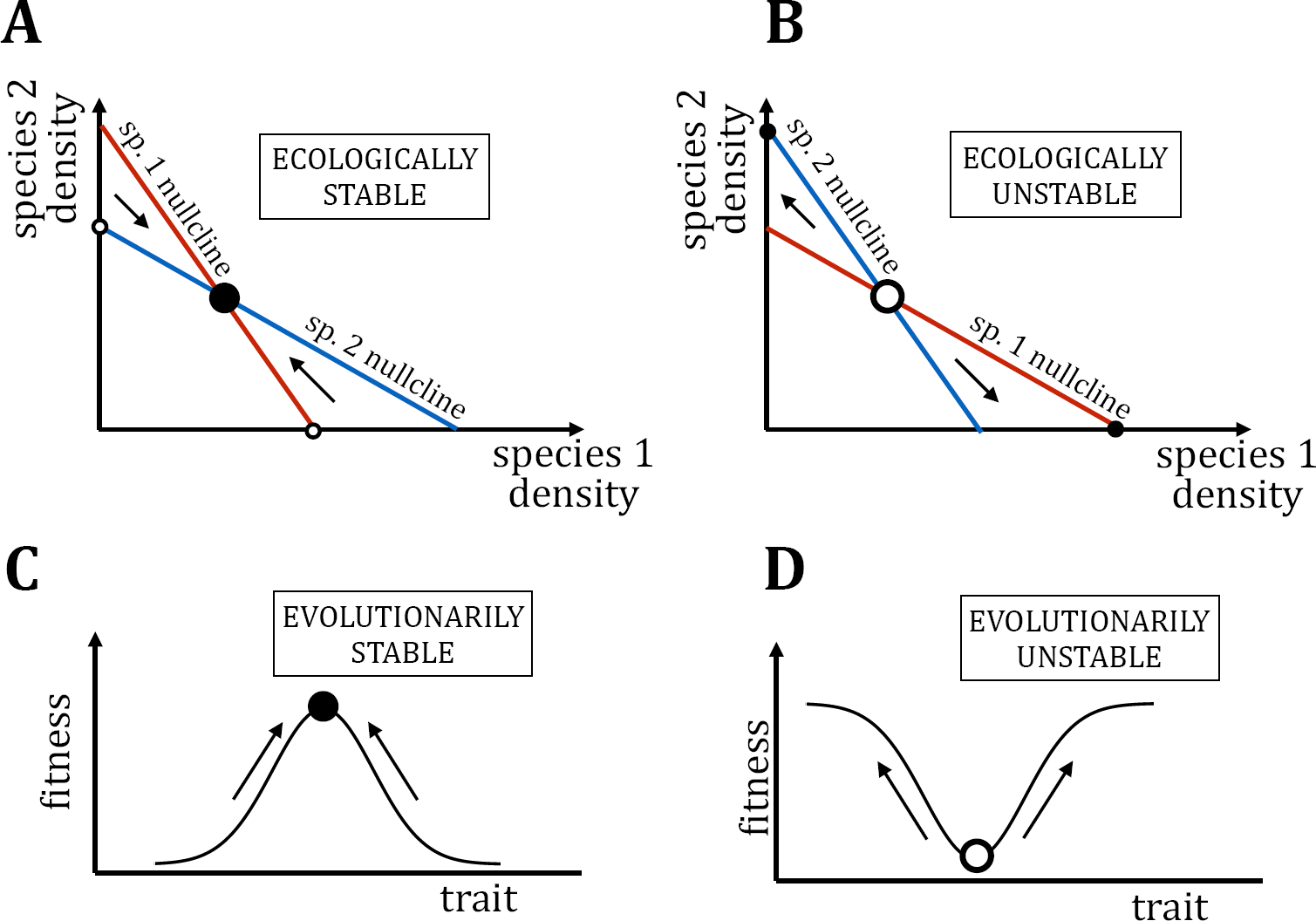
Schematics of stable and unstable equilibria in a classic two competing species ecological model (A, B) and an evolutionary model (C, D). When intraspecific competition is greater than interspecific competition, the coexistence equilibrium between two competing species is stable (A) and otherwise unstable (B). With frequency independent selection, equilibria at local fitness maxima are stable (C) and at local fitness mimima are unstable (D).

We assume that one of the two competing species (species 1) has a quantitative trait that is subject to frequency independent selection. Under the Lande (Lande 1976) framework, this implies that a trait equilibrium occurs when the fitness gradient is zero (see Equation 2) and is evolutionarily (un)stable when fitness is locally (minimized) maximized (Figure 3). We examine two different types of traits and show that in one case, eco-evolutionary feedbacks are stabilizing for slow and fast evolution, while in another they are destabilizing. Through these two examples, we show how purely qualitative information (signs of effects) can be used to determine the role of eco-evolutionary feedbacks on stability.

### Competition Model I: Eco-evolutionary feedbacks are stabilizing

Both competing species experience intra- and interspecific competition, so that growth rates are negatively related to population densities. Therefore, the ecological Jacobian matrix is

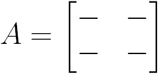

with the negative effects of intra- and interspecific competition captured in the signs of the diagonal and off diagonal elements, respectively.

In the first model, we assume that the evolving trait in species 1 exhibits a trade off in the ability of individuals to compete with conspecifics versus heterospecifics. Following Vasseur et al. (2011), this trait reduces the antagonistic effects of species 2 on species 1, increases the antagonistic effects of species 1 on species 2, and increases intraspecific antagonism (Figure 4A). Traits such as body size or aggressiveness fall into this category. For example, aggressiveness in social spiders (*Anerlosimus studious*) leads to increased competitive ability against heterospecifics but a reduction in resource-use efficiency, increasing intraspecific competition (Pruitt and Riechert 2009).

**FIGURE 4.**
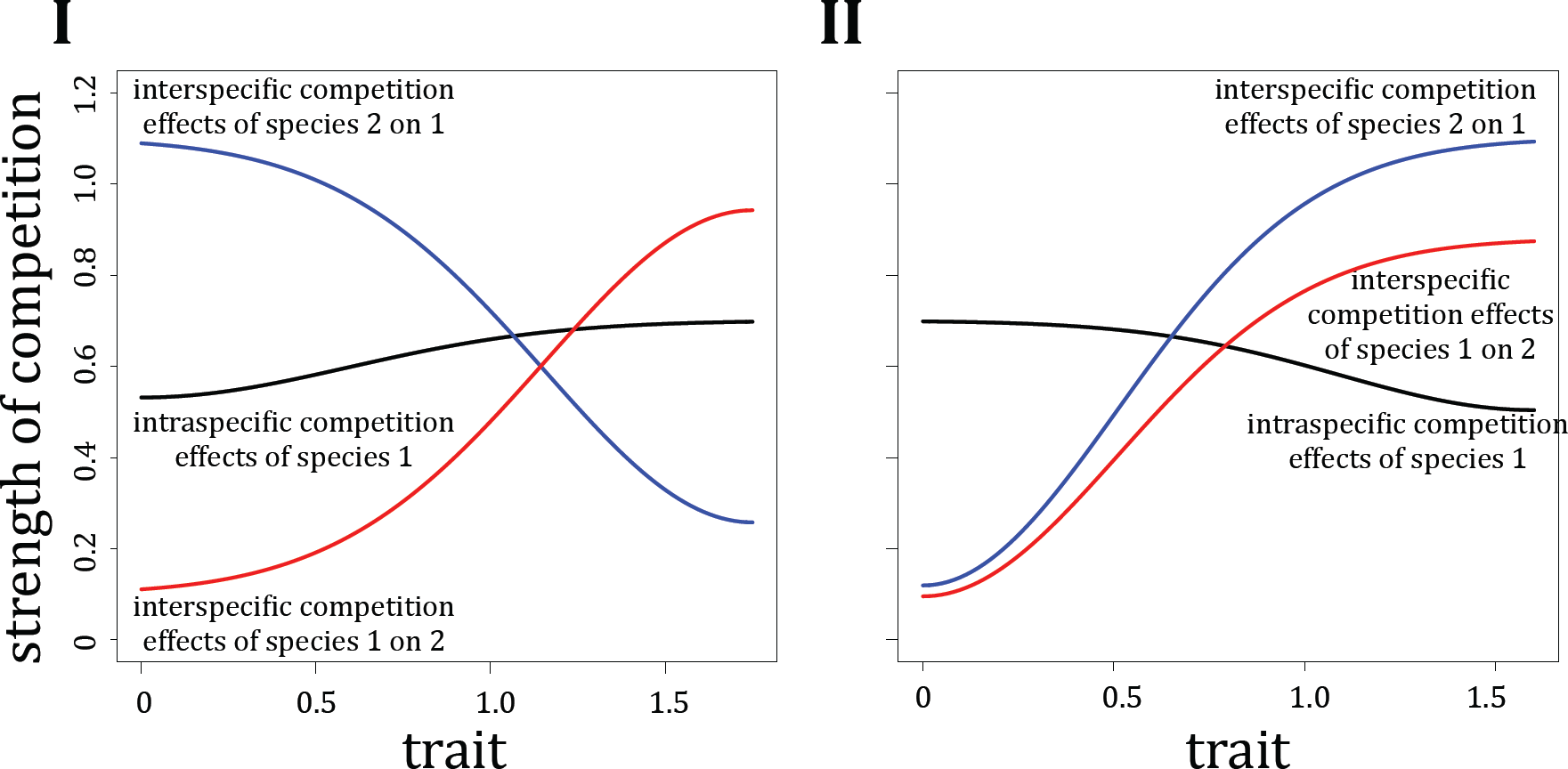
Intra- and interspecific competition for traits in competition models I and II. In model I, increasing the trait value increases intraspecific competition within species 1 as well as interspecific competition effects of species 1 on 2 while decreasing interspecific competition effects of species 2 on 1. In model II, increasing the trait values increases interspecific competition effects between species 1 and 2 while decreasing intraspecific competition effects within species 1. For details of model, see Appendix A4.

This qualitative information about the type of trait determines the signs of *b* and *c* vectors in the Jacobian matrix. In particular, there are four important relationships between demography and selection that determine the effects of eco-evolutionary feedbacks on stability. First, increased trait values reduce the fitness of species 2. Second, increased density of species 1 selects for decreasing the trait. Third, increased density of species 2 selects for increasing the trait. Finally, consistent with Vasseur et al. (2011), since selection is frequency independent, the fitness gradient equals zero at a trait equilibrium (Lande (1976), Figure 3). Altogether, these four statements determine the signs of vectors *b* and *c* in the Jacobian matrix at an equilibrium.

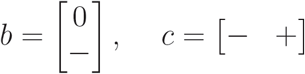

When the evolutionary dynamics are slow relative to the ecological dynamics, the eco-evolutionary feedback can stabilize an evolutionarily unstable equilibrium. Indeed, the sign of the evo-eco-evo pathway is given by

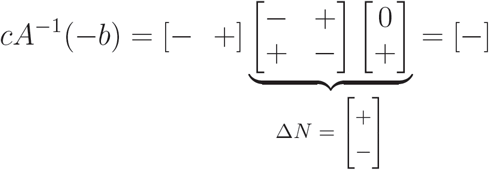

In the first step of this pathway, a small increase in the trait leads to an increase in the density of species 1 and decrease of species 2 (from Δ*N*). These population density changes result in a selective pressure for decreased trait values, which counteracts the perturbation. This creates a negative feedback, which has stabilizing effects (corresponding to the bottom half of Figure 2B) and can stabilize an equilibrium that is evolutionarily unstable (*d >* 0).

Similarly, when the evolutionary dynamics are fast relative to the ecological dynamics, the eco-evolutionary feedback can stabilize an ecologically unstable equilibrium. The stabilizing feedback comes from the eco-evo-eco pathway, given by

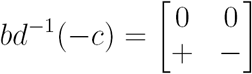

Analogous to the community matrix of direct ecological effects, this 2*×*2 matrix represents the indirect effect population densities have on each other’s growth via an indirect pathway. In general, it is difficult to determine the effects of adding matrices on the stability. However, for 2*×*2 matrices, the stability modulus is negative if the trace is negative and the determinant is positive. Adding the eco-evolutionary feedback term to matrix *A* has two effects: it increases the product of the diagonal terms and it decreases the product of the off-diagonal terms. This makes the determinant of *A* + *bd*^*−1*^(*−c*) more positive and the trace more negative, which are both stabilizing effects.

To summarize, the eco-evo-eco and evo-eco-evo feedbacks are both stabilizing. But when do they actually result in a stable equilibrium? To address this question, we compared our general theory to the specific two-species competition differential equation model from Vasseur et al. (2011). In this model, we examined stability from slow to fast evolution for a range of two parameters: (1) the strength of intraspecific competition of the non-evolving species, which is independent of the trait, and (2) the distance between the trait values that are optimal for intra- and interspecific competition of the evolving species (trait trade off) (see Appendix A4 for more details; Figure 5). We choose these two parameters because of their targeted effect on ecological stability and evolutionary stability, separately. Increasing the strength of intraspecific competition of the non-evolving species increases ecological stability (decreases *s*(*A*)) with little effect on evolutionary stability. Increasing the trade off decreases evolutionary stability (decreases *d*) with little effect on ecological stability (Figure 5). Hence, by varying these parameters, we can test for the stabilizing effects of eco-evolutionary feedbacks on four types of equilibria: (1) ecologically and evolutionarily stable, (2) ecologically stable and evolutionarily unstable, (3) ecologically unstable and evolutionarily stable, and (4) ecologically and evolutionarily unstable.

**FIGURE 5.**
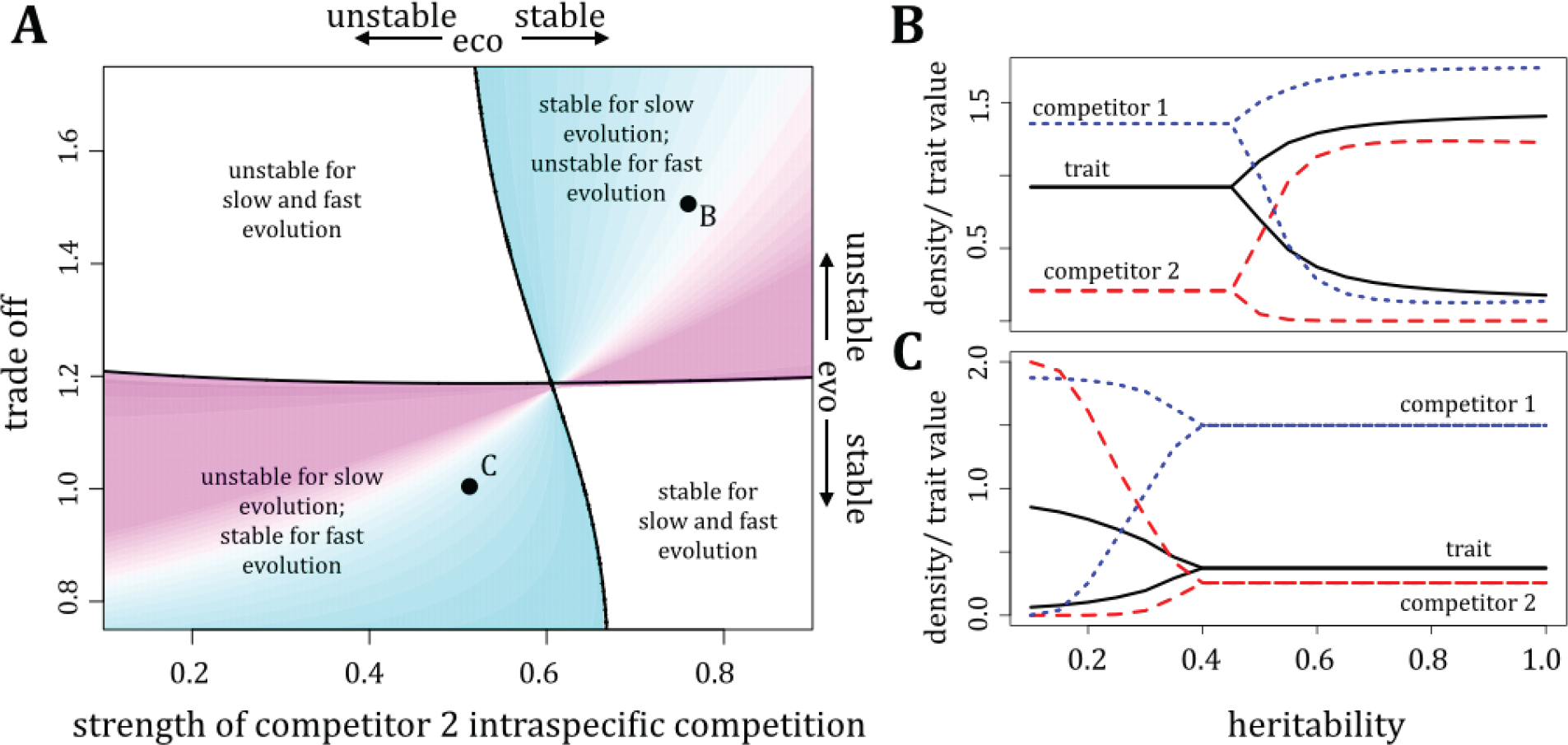
Ecological stability, evolutionary stability and overall stability for competing species in model I. In (A), the x-axis is the strength of intraspecific competition of the non-evolving species and the y-axis is the strength of the trade off between intra- and interspecific competition. The vertical and horizontal black curves correspond to *s*(*A*) = 0 and *d* = 0, separating ecological stability and evolutionary stability, respectively. Each resulting region is labeled by whether the equilibria are overall stable for slow and fast evolution. In the top right and bottom left, there is a switch in stability at a critical rate of evolution. The color gradient corresponds to when the switch in stability occurs, with light blue representing the slowest critical rate and pink representing the fastest. (B) and (C) show the long-term minimum and maximum densities and trait values for varying heritabilities, which determines the relative rate of evolution. The parameters correspond to points B and C in (A).

Our numerical results match predictions from our theory on the effects of eco-evolutionary feedbacks for slow and fast evolution. First, since feedbacks are stabilizing, if an equilibrium is both ecologically stable (*s*(*A*) *<* 0) and evolutionarily stable (*d <* 0), then the equilibrium is stable for slow and fast evolution (bottom right in Figure 5). We note that our theory applies when evolution is slow or fast but in general, does not determine stability at intermediate rates (see, however, Appendix A3 for a special exception). By incrementally varying the heritability, which determines the relative rate of evolution to ecological dynamics in the model, we found that the equilibria were always stable for intermediate rates of evolution.

Second, if an equilibrium is ecologically stable and evolutionarily unstable (*d >* 0), then the evo-eco-evo feedbacks can stabilize this equilibrium when evolution is slow (iv in Figure 2B, C and top right in 5A). For faster evolutionary rates, our theory indicates that these equilibria will be unstable, as the unstable direct evolutionary effects become increasingly important (Figure 1). In 5A (top right), we show the critical relative rate at which this loss of stability occurs. An equilibrium that is more ecologically stable (smaller *s*(*A*)) has a larger critical relative rate to lose stability than equilibria that are less ecologically stable (horizontal transects in 5A top right).

Third, if an equilibrium is ecologically unstable and evolutionarily stable, then the eco-evo-eco feedbacks can stabilize the equilibrium when evolution is fast (bottom left in 5). For slower evolutionary rates, our theory indicates that these equilibria will be unstable, as the unstable direct ecological effects become increasingly important (Figure 1). In 5A (bottom left), we show the critical relative rate at which this change in stability occurs. An equilibrium that is more evolutionarily stable (smaller *d*) has a lower critical relative rate that changes stability (vertical transects in 5A bottom left).

Finally, if an equilibrium is ecologically and evolutionarily unstable, then it is unstable for slow and fast evolution. In our numerical solutions, these equilibria were unstable for all relative rates of evolution.

### Competition Model II: Eco-evolutionary feedbacks are destabilizing

Different traits may lead to different relationships between demography and selection, thereby affecting the role of eco-evolutionary feedbacks on stability. Suppose now that the evolving competitor (species 1) has a trait that increases the antagonistic interactions between species 1 and 2, while decreasing intraspecific antagonism (Figure 4B). An example of such a trait is when time spent on antagonistic interactions with interspecific competitors reduces time spent on antagonistic interactions with intraspecific competitors. More specifically, in territorial organisms, competition with heterospecifics for space can reduce individual range overlap with conspecifics (Hoi et al. 1991).

There are two key differences from traits in model I: increased density of species 1 selects for higher trait values while increases in species 2 select for lower trait values. These differences are captured in the signs of vectors *b* and *c*

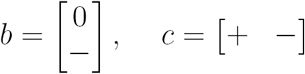

The feedback term for slow evolution (evo-eco-evo pathway) is

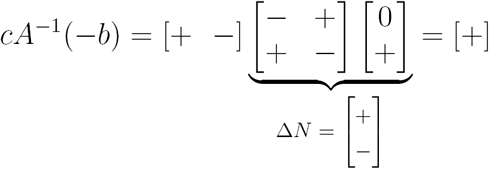

The first step of this pathway is the same as in model 1: a small increase in the trait leads to an increase in the density of species 1 and decrease of species 2 (from Δ*N*). However, these population density changes result in a selective pressure to further increase the trait values, which magnifies the perturbation. This creates a positive feedback, which has destabilizing effects (top half of Figure 2B). Thus, even if the equilibrium is ecologically and evolutionarily stable, it can be overall unstable due to these destabilizing evo-eco-evo feedbacks (iii in Figure 2B,C).

When evolution is fast, the critical eco-evolutionary feedback is via the eco-evo-eco pathway given by

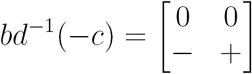

which is destabilizing because adding *bd*^*−1*^(*−c*) to *A* decreases the determinant and increases the trace.

To summarize, the important eco-evolutionary feedbacks for slow and fast evolution are both destabilizing, which is the opposite of that found for traits in model I. We tested this general theory with a model modified from Vasseur et al. (2011) to reflect traits of this type (see Appendix A4). Consistent with our theory, we find that eco-evolutionary feedbacks are often sufficiently destabilizing such that equilibria with stable ecological and evolutionary subsystems are overall unstable (Appendix A4).

## EFFECTS OF CORRELATED MULTI-TRAIT EVOLUTION ON STABILITY

Previous work has shown that in purely evolutionary models when a single species is evolving in multiple quantitative traits that contribute independently to fitness, correlations between these traits will not qualitatively affect evolutionary stability (Lande 1979). Correlations can, however, affect the degree of stability, and slow down or speed up the return to a stable equilibrium, following a perturbation. Here, we ask what is the role of correlations on stability in light of eco-evolutionary feedbacks.

First, we ask whether correlations can qualitatively change stability when multiple quantitative traits of a single species within the community evolve (using equation (3)). For simplicity, we assume that all the traits have equal genetic variances so that the genetic covariance in equation (3) can be written as *v*_*mj*_ = *vρ_mj_*, where *v* is the constant genetic variance and *ρ*_*mj*_ is the genetic correlation between traits *m* and *j*. Then, if we let *ϵ* = *v*, we can express matrices *C* and *D* in the Jacobian (4) as 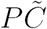 and 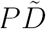, where *P* is the genetic correlation matrix with elements *ρ*_*mj*_,

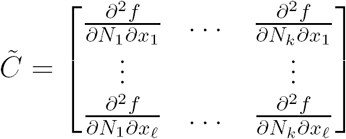

and 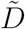 is a diagonal matrix with elements 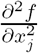, where *f* is the fitness of the evolving species. When evolution is slow, the conditions for overall stability are ecological stability (*s*(*A*) *<* 0; unaffected by correlations) and 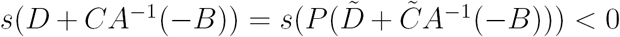. As the correlation matrix *P* can transform the evo-eco-evo feedback, correlations can change stability. On the other hand, when evolution is fast, the conditions for overall stability are evolutionary stability 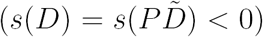 and 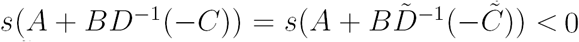. As Lande (1979) has shown, the sign of 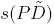 is unaffected by correlation matrix *P*, correlations between multiple traits do not qualitatively affect stability when evolution is sufficiently fast.

To test for the effects of correlations on stability with eco-evolutionary feedbacks, we apply our results to a three-species Lotka-Volterra food chain model, in which two traits of the herbivore evolve, one that determines its ability to consume the basal species and another that influences its defense against the top predator. We assume that there is a trade off between resource consumption and intraspecific competition, and a trade off between defense and intraspecific competition. To model these trade offs, we use modified versions of the functions from Schreiber et al. (2011) and Vasseur et al. (2011). While we assume that these traits act independently to determine the interaction strengths with the basal species and top predator, we allow for the two traits to be genetically correlated (see Appendix A5 for more details). While these model assumptions determine the sign structure of matrices *A, B* and 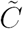, this sign structure is not sufficient to indicate whether the eco-evo-eco nor the evo-eco-evo feedback is stabilizing or destabilizing (Appendix A5). Consequently, we explore this model numerically.

We plot contours of the stability modulus of the Jacobian (4) for varying genetic variances (evolutionary rates) and correlations between the two traits (Figure 6A). We observe both the qualitative effects (transition between stable and unstable; thick black curve in Figure 6A) and quantitative effects (degree of stability; gray contours in Figure 6A). From classic Lotka-Volterra food chain theory, when an equilibrium supporting all three species exists, it is ecologically stable (*s*(*A*) *<* 0; (Hofbauer and Sigmund 1998)). We choose parameters so that the equilibrium is always evolutionarily unstable and find a U-shaped relationship between the effect of the genetic variance and correlation on stability (Figure 6A). When evolution is slow, evo-eco-evo feedbacks always stabilize the equilibrium for all correlations (Figure 6A). As predicted by our theory, correlations affect the stability modulus of the evo-eco-evo feedback and, consequently, the degree of stability. Correlations, whether positive or negative, increase the stability modulus and thereby decrease the degree of stability for slow evolution (Figure 6Bi). As our theory predicts for fast evolution, correlations do not affect the stability modulus with the eco-evo-eco feedback and these feedbacks do not stabilize the equilibrium (Figure 6A, Bii). However, correlations, whether positive or negative, decrease the stability modulus of the evolutionary dynamics and thereby decrease the degree of instability (Figure 6Bii). Note that in Figure 6A, for extremely negatively or positively correlated traits, the equilibrium is stable, but this diminishes for even faster evolutionary rates. Finally, for intermediate rates of evolution, correlations qualitatively change stability: only when the two traits are sufficiently positively or negatively correlated is the equilibrium overall stable (Figure 6C).

**FIGURE 6.**
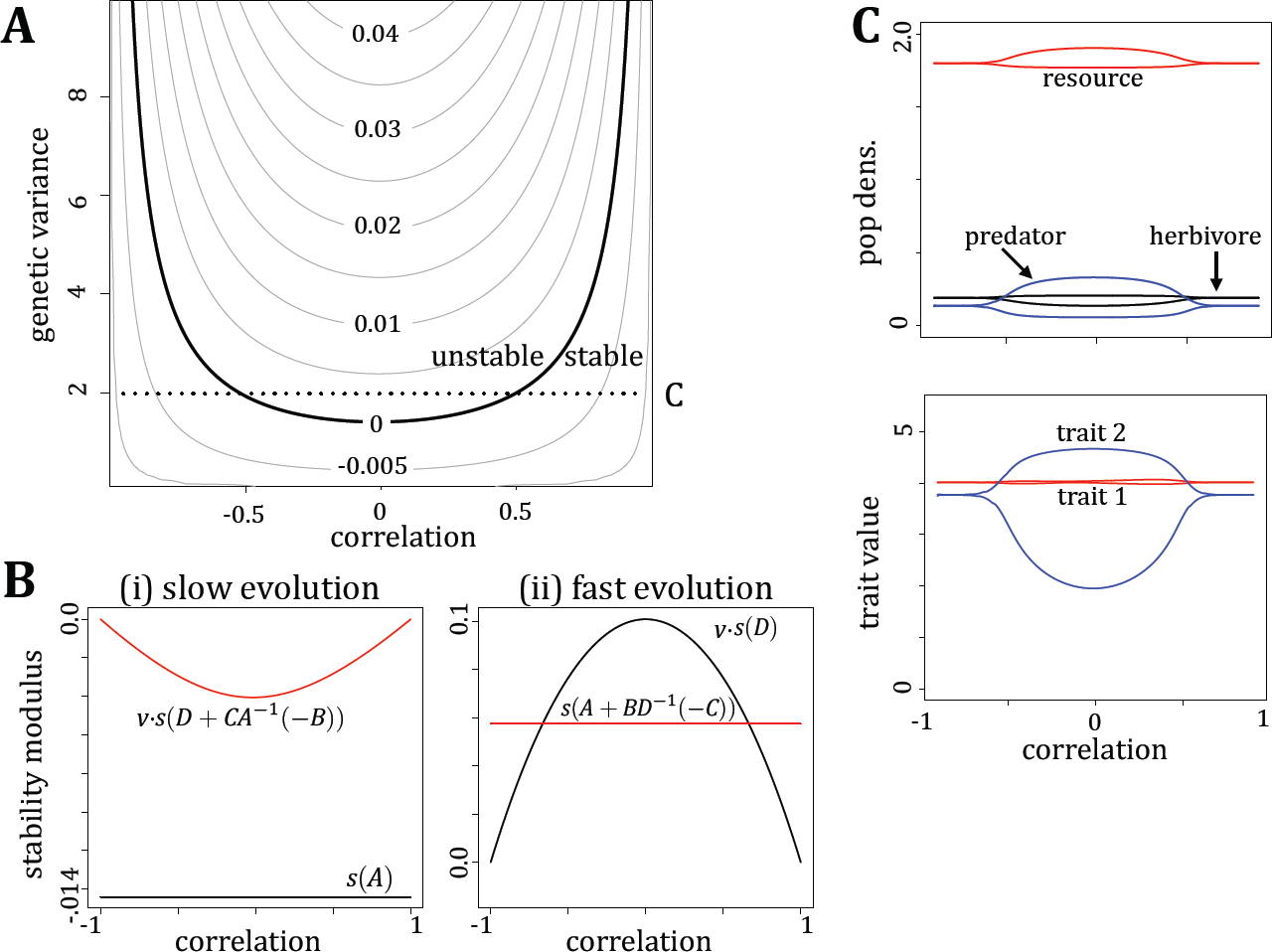
The effects of trait correlations on stability of food chains. (A) shows contours of the stability modulus for varying genetic variances (rates of evolution) and correlations. (B) plots the two relevant stability moduli from our stability conditions for slow (i; *s*(*A*) and *s*(*D* + *CA*^*−1*^(*−B*)); *v* = 0.099) and fast (ii; *s*(*D*) and *s*(*A* + *BD*^*−1*^(*−C*)); *v* = 9.9) evolution with the eco-evolutionary feedback condition in red. (C) plots the minimum and maximum population densities and trait values at the eco-evolutionary attractor for a fixed rate of evolution (corresponding to the horizontal dotted line in (A)) and varying correlations.

## DISCUSSION

### Implications of our results

Existing empirical and theoretical studies have demonstrated that eco-evolutionary feedbacks can have substantial effects on community stability, but this has largely focused on specific ecological modules (Abrams and Matsuda 1997; Becks et al. 2010; Cortez and Ellner 2010; Vasseur et al. 2011; Schreiber et al. 2011; Steiner and Masse 2013; Patel and Schreiber 2015; Cortez 2016). Our analysis unifies these existing studies and provides a general theory for mechanistically understanding how eco-evolutionary feedbacks impact stability, which can be applied to model communities with any number of species and evolving traits. In particular, we provide conditions for stability for when evolution is fast and slow relative to ecology. By interpreting these conditions, we draw inferences on when and how direct ecological, direct evolutionary and indirect eco-evolutionary feedbacks impact stability and how these impacts depend on the relative time scales of the ecological and evolutionary processes.

Recently, Hendry (2016) highlighted that understanding the effects of the relative rate of evolutionary processes to ecological processes and the role of direct and indirect effects on community stability is a fundamental problem in eco-evolutionary dynamics. Our work makes several advances on this problem. The conditions for stability when evolution is slow relative to ecology are that the direct ecological effects must be stable and the sum of the direct evolutionary effects with the indirect evo-eco-evo feedbacks must be stable. In contrast, when evolution is fast, the direct evolutionary effects must be stable and the sum of the direct ecological effects with the indirect eco-evo-eco feedbacks must be stable. As we describe in the main text, these results have a graphical interpretation as a two-step response to perturbations in systems with a separation of time scales. Importantly, these conditions highlight that on one hand, eco-evolutionary feedbacks can destabilize an equilibrium that is ecologically and evolutionarily stable and on the other hand, they can stabilize an equilibrium that is unstable in the slower dynamic.

Our theory indicates that an equilibrium may not be stable for all evolutionary time scales, i.e., system stability can depend on the relative rate of evolution to ecology. In particular, it may be stable for slow evolution and unstable with fast evolution or vice versa. We can draw two general conclusions that are true for all types of species interactions in this framework. First, if an equilibrium is evolutionarily unstable but stabilized by the slow evo-eco-evo feedback, then the equilibrium will destabilize when evolution occurs sufficiently quickly. For example, in the first model of traits for two competing species, this destabilization occurs despite the eco-evolutionary feedbacks having a stabilizing effect for slow and fast evolution (Figure 5). Second, if an equilibrium is ecologically unstable but stabilized by the fast eco-evo-eco feedback, then it will lose its stability with slow evolution. These stability changes for varying rates of evolution have been observed theoretically in predator-prey (Abrams and Matsuda 1997; Cortez 2016), competition (Vasseur et al. 2011), and intraguid predation communities (Patel and Schreiber 2015) as well as suggested experimentally in predator-prey communities (Becks et al. 2010). Our general results suggest that one cause for the loss or gain of stability with varying rates of evolution observed in these studies may be the switch in the dynamics that dominates the fast response.

Our three examples show that the main results can be applied to communities with different types of species interactions and evolutionary constraints. Applying our results to two species competition, we demonstrate how qualitative information about the relationships between traits and population growth sometimes is sufficient to infer the effects of eco-evolutionary feedbacks on stability. Importantly, qualitative differences in these relationships can alter whether the feedback is stabilizing or destabilizing. For a trait that increases the competitive ability of the evolving species against heterospecifics at the cost of increasing competition amongst conspecifics (competition model I), eco-evolutionary feedbacks are stabilizing. In contrast, when the trait increases interference competition between heterospecifics (competition model II) while reducing competition amongst conspecifics, eco-evolutionary feedbacks are destabilizing. In both cases, in the evo-eco-evo feedback, a small positive perturbation in the trait leads to an increase and decrease in the density of species 1 and 2, respectively. In the first case, these population density changes drive a selection pressure that opposes the perturbation. In the second case, these population density changes drive selection pressures that amplify the initial trait perturbation.

When a single species evolves in multiple quantitative traits, classic evolutionary theory asserts that correlations cannot qualitatively impact evolutionary stability (Lande 1979). These correlations can, however, alter the trajectories of how traits approach (i.e. stabilizing selection) or diverge (e.g. disruptive selection) from an equilibrium. When evolution is sufficiently fast relative to ecology, traits evolve very quickly to the equilibrium relative to the ecological process and so the trajectories traits take to get to the equilibrium (and hence, correlations) have little effect on the overall eco-evolutionary feedbacks. However, as the relative rate of evolution becomes more commensurate or slower than the ecological dynamics, the traits no longer reach their equilibrium before eliciting an ecological response. Hence, the correlation effects on trait trajectories become important in the eco-evolutionary feedbacks. Importantly, in this way, stability depends critically on correlations when evolution occurs at slow or intermediate rates relative to ecology.

### Comments on our framework

We illustrated our results with Lande’s quantitative genetics framework (Lande 1976). Our results are also applicable to two additional notable existing frameworks of modeling evolutionary dynamics: the explicit multi-locus evolutionary framework and the adaptive dynamics framework. Hence, our results can be applied to many distinct descriptions of the evolutionary process.

The multi-locus modeling framework describes the evolutionary process by characterizing fitness differences amongst explicit genotypes in the population and tracking genotypic frequencies as the evolutionary variable (Doebeli 1997; Yamaguchi et al. 2011; Yamamichi and Ellner 2016). In this framework, the genotypic frequencies are the population-level quantitative values of interest. When fitness differences between genotypes are small, evolution may occur slowly relative to ecological population density changes. On the other hand, if population growth is constrained, then ecological dynamics may occur slowly relative to evolutionary genotype frequency changes.

In the adaptive dynamics framework, populations are assumed to be monomorphic and clonal, with evolution occurring due to mutations of small effect. Since it is assumed that mutations are rare, the evolutionary dynamics are assumed to be much slower than the ecological dynamics. Hence, our mathematical results for slow evolution can also be applied to an adaptive dynamics framework for studying eco-evolutionary dynamics. In Appendix S3, we describe in more detail the interpretations of the stability conditions in an adaptive dynamics framework.

Finally, time scale separations may also occur among or within behavioral, ecological, or ecosystem processes relevant to the system dynamics. For all of these scenarios, the stability conditions presented here can still be applied. For example, our conditions can be used to understand stability in models with behavioral or plastic traits, which are often thought to change very quickly relative to ecological processes (e.g. Abrams 1992; Ma et al. 2003), to purely ecological models, in which a subset of species naturally reproduces and dies at a faster rate than another set of species (e.g. Muratori and Rinaldi 1992), or to combinations of these cases as in Takimoto et al. (2009). Importantly, the responses of a system to perturbations may depend on which subset of processes occurs slowly relative to the remaining set of processes (Takimoto et al. 2009).

### Future directions

Our analysis shows that for sufficiently slow evolutionary rates, the evo-eco-evo feedback is critical, while for sufficiently fast evolutionary rates, the eco-evo-eco feedback is critical. What does this mean for intermediate rates of evolution? Except for the special case in which the slow dynamics are one-dimensional (see Appendix A2), in general we cannot make stability conclusions for intermediate rates of evolution based on our understanding of fast and slow evolution. For intermediate rates of evolution, additional eco-evolutionary interactions not accounted for in the feedback pathways given for slow and fast evolution will play a role and can alter stability. In particular, even if an equilibrium is stable for slow and fast evolution, it may be unstable for intermediate rates of evolution (for example see Cortez (2016) Figure 4E). Simple and general analytical results for partitioning eco-evolutionary feedback effects for intermediate rates are limited. Further development in this area may help to understand the role of eco-evolutionary feedbacks on stability and bridge the slow and fast limits.

Through our competition examples, we showed that purely qualitative information about the relationships between traits and population growth (signs of the matrices) were sufficient for predicting whether eco-evolutionary feedbacks were stabilizing or destabilizing. In most cases of more complex models, sign structure alone will not be sufficient to infer the effects of eco-evolutionary feedbacks as in the two species competition cases considered here. This is consistent with the pattern May (1973) noted when he used sign structure of purely ecological community matrices to determine stability. Hence, for more complex models, our theory is still applicable, when we can obtain quantitative information about these eco-evolutionary relationships, i.e., the Jacobian elements.

One way in which we can obtain this information is to use empirical time series data to parameterize a model and then estimate the Jacobian (Deyle et al. 2016). From this, we can observe whether there is an evident time scale separation, and, if so, apply our method to partition out the influence of pure ecology, pure evolution, and eco-evolutionary feedbacks on stability. Alternatively, if the time scales of ecology and evolution are similar, then we can ask how stability would be different if one process was sped up or slowed down to occur much more quickly than the other. By applying these methods to empirical systems, we might be able to identify the relative contributions of ecology, evolution, and eco-evolutionary feedbacks on community stability or instability.

Finally, in this work, we focus particularly on the effects of eco-evolutionary feedbacks on community stability, which determines how a community at equilibrium will respond to small yet potentially frequent perturbations (Schreiber 2006). While stability is a fundamental property of communities, there are many other properties of interest to community ecologists, including the persistence of species, resilience to large perturbations, and robustness (Hutson and Schmitt 1992; Schreiber 2000; Meszéna et al. 2006; Barabás et al. 2012; Barabás and D’Andrea 2016). Developing a general framework for identifying the effects of feedbacks on these and other critical properties of communities is an important avenue for future research to build a broader understanding of how communities will respond in the face of new ecological and evolutionary pressures.

## ACKNOWLEDGEMENTS

We thank Thomas Schoener, Jaime Ashander, Axel Saenz, Casey terHorst, Peter Zee and Sam Fleischer for useful discussion on this topic and comments on the manuscript. Financial support by the U.S. National Science Foundation Grant DMS-1313418 to SJS, the American Association of University Women Dissertation Fellowship to SP and the Austrian Science Fund (FWF) Grant P25188-N25 to Reinhard Burger at the University of Vienna is gratefully acknowledged.

## A1. PROOF OF GENERAL STABILITY CONDITION

With *k* interacting species and *l* traits evolving, the model is

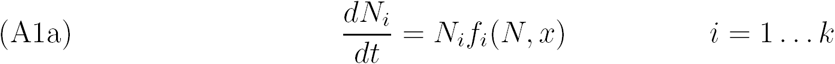

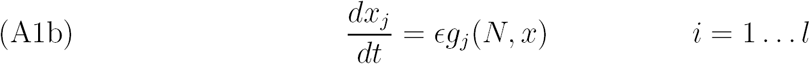

For a matrix *M*, we denote the stability modulus, i.e. the largest real part of the eigenvalues, as *s*(*M*). The Jacobian matrix of (A1) at an equilibrium is

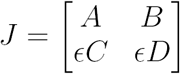

with sub matrices *A, B, C, D*, as described in the main text.

**Theorem A1.** *If s*(*A*) *<* 0 *and s*(*CA*^*−1*^(*−B*) + *D*) *<* 0 *at a hyperbolic equilibrium of* (A1), *then the equilibrium is stable, for sufficiently small ϵ <* 0.

*Proof.* To find all the eigenvalues of *J*, we must find solutions to the eigenvalue equation 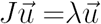. Following a standard “balancing procedure” singular perturbation method for finding solutions to algebraic equations, we will treat *ϵ* as a perturbation and use the perturbation expansions

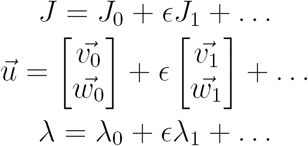

where

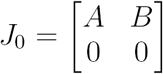

and

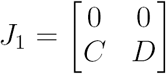

Applying these expansions to the eigenvalue equation and equating coefficients of *ϵ*^0^ and *ϵ*^1^ yields

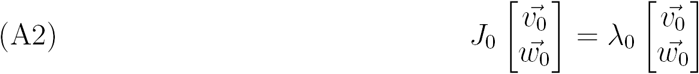

and

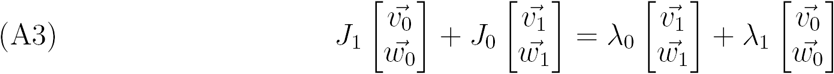

From (A2), we get the following two equations must hold

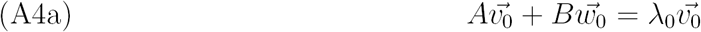

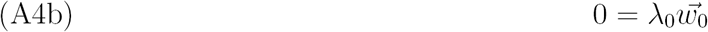

From (A3), we get that the following two equations must hold

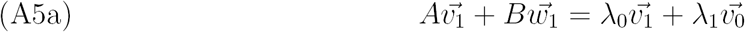

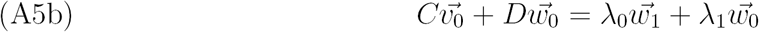

Eigenvalues will fall into one of two cases: (i) *λ*_0_ ≠ 0 or (ii) *λ*_0_ = 0. In case (i), 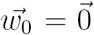 and *λ*_0_ are eigenvalues of *A*, which follows from (A4). Since *s*(*A*) *<* 0, these eigenvalues have negative real part, and will remain this way with sufficiently small perturbations of *ϵ*.

In case (ii), *λ*_0_ = 0 and we must look at higher order terms (*λ*_1_) to determine whether these eigenvalues have negative real part. From (A4a), we see that 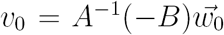. Applying this relationship to (A5b), we see that *λ*_1_ must satisfy 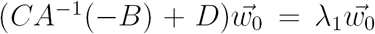 and hence, must be an eigenvalue of *CA*^*−1*^(−*B*) + *D*. If *s*(*CA*^*−1*^(−*B*) + *D*) *<* 0, then the higher order terms will all have negative real part. □

The stability conditions for equilibria in models with fast evolution are identical. Since the single trait models are a subset of multi-trait models, these results apply to the single trait models as well.

### A2. PROOF OF STABILITY INVARIANCE WITH TIME SCALE PARAMETER IN A SPECIAL CASE

Here we consider a special case of ecological dynamics coupled to one evolving trait, in which we can determine stability for intermediate rates of evolution from our conditions for stability for slow evolution. With *k* interacting species and one evolving trait, model (1) in the main text simplifies to

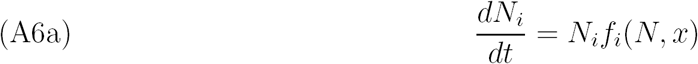

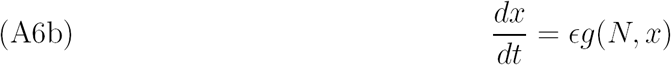

The following theorem shows that in this special case, if an equilibrium is ecologically stable and overall unstable for slow evolution, then it is unstable for all rates of evolution.

**Theorem A1.** *If s*(*A*) *<* 0 *and s*(*cA*^*−1*^(*−B*) + *d*) *>* 0 *at a hyperbolic equilibrium of* (A6), *then s*(*J*_*ϵ*_) *>* 0 *for all ϵ <* 0, *where A, B, c and d are defined in the main text and J_ϵ_ is (5) with fixed ϵ.*

*Proof.* First, assume the dimensions of the ecological dynamics (*k*) are even. Ecological stability (*s*(*A*) *<* 0) implies that det(*A*) *>* 0. If *s*(*CA*^*−1*^(*−B*) + *D*) *>* 0, then the equilibrium is overall unstable for sufficiently small 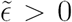. Hence, det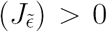. Signs of determinants are invariant for all *ϵ >* 0, and so, det(*Jϵ*) *>* 0 for all *ϵ >* 0, which implies the equilibrium is unstable for all *ϵ >* 0. The argument holds for when ecological dimensions are odd by reversing all the inequalities. □

### A3. COMPARISON TO ADAPTIVE DYNAMICS APPROACH

Here we discuss the connections between the stability results developed in this study and the stability results from the theory of adaptive dynamics (Marrow et al. 1996; Geritz et al. 1998). The relations between the conditions are summarized in Table A3. Throughout we will only focus on systems in the slow evolution limit and when selection is frequency independent. We also assume that the ecological subsystem is stable.

In the adaptive dynamics approach, the ecological dynamics of the system are assumed to be much faster than the evolutionary dynamics. Specifically, the approach assumes monomorphic populations undergoing clonal reproduction, that all populations converge to an ecological equilibrium before mutant offspring arise, and that mutations are of small effect, implying that the trait values of mutant offspring are very close to the mean trait value. The equation governing the evolution of the mean trait value (*x*_*i*_) for species *i* under frequency independent selection is

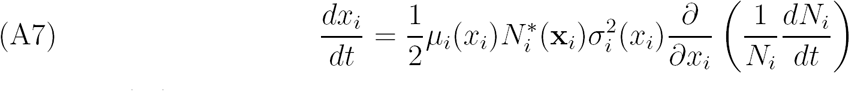

where 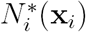 is the equilibrium density of species *i* when the mean traits are **x***_i_*, *μ*_*i*_(*x*_*i*_) is the probability of mutation, and *σ*_*i*_(*x*_*i*_) is the variance of the mutation distribution.

Before comparing the results of this study to those from the theory of adaptive dynamics, we define a few terms from the adaptive dynamics framework. An *Evolutionary Singular Strategy* (ESS) is a trait value at an equilibrium point. A *Convergent Stable Strategy* (CVS) is an ESS that is an evolutionary attractor, i.e., a trait value at an equilibrium that is stable in the slow evolution limit. An ESS that is not a CVS is an evolutionary repellor. An *evolutionary branching point* is a CVS where 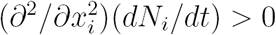 for some *i*. In our framework, a CVS is an equilibrium point where the matrix *D* + *CA*^*−1*^(−*B*) has all eigenvalues with negative real part, i.e., *s*(*D* > *CA*^*−1*^(−*B*)) *<* 0, and an evolutionary branching point is an equilibrium point where at least one of the diagonal entries of *D* is positive. Note that evolutionary branching in one species can induce evolutionary branching in other species.

First consider systems with a single trait, i.e., one-dimensional trait dynamics. In this case, the CVS condition is *d* + *cA*^*−1*^(−*b*) *<* 0. This means that an equilibrium point is a CVS either when (i) the evolutionary subsystem is unstable (*d >* 0) and the eco-evolutionary feedback stabilizes the system or (ii) the evolutionary subsystem is stable (*d <* 0) and the eco-evolutionary feedback does not destabilize the system. The evolutionary branching condition for systems with a single trait is *d >* 0. This means that evolutionary branching occurs for every CVS where the evolutionary subsystem is unstable and evolutionary branching does not occur when the evolutionary subsystem is stable. Hence, in the case where the evolutionary subsystem is unstable, the stabilization caused by the eco-evolutionary feedback allows for evolutionary branching to occur. Finally, note that if the evolutionary subsystem is unstable and the eco-evolutionary feedback does not stabilize the equilibrium, then the equilibrium trait value is a repellor.

Now consider systems with multiple traits, i.e., multiple evolving species or one species with multiple evolving traits. As with the previous case, an equilibrium is a CVS either when (i) the evolutionary subsystem is unstable and the eco-evolutionary feedback stabilizes the system or (ii) the evolutionary subsystem is stable and the eco-evolutionary feedback does not destabilize the system. There are two important points to make about evolutionary branching in these systems. First, note that because the evolutionary branching requires at least one diagonal entry of *D* to be positive, instability of the evolutionary subsystem does not guarantee evolutionary branching. For example, if *D* is 2x2 with negative diagonal entries and a negative determinant and the eco-evolutionary feedback stabilizes the equilibrium, then the equilibrium will be an CVS, but not an evolutionary branching point. However, if one of the diagonal entries of *D* is positive and the eco-evolutionary feedback stabilizes the equilibrium, then the equilibrium point will be an evolutionary branching point. Second, if the evolutionary subsystem is stable, then evolutionary branching can occur, provided at least one diagonal entry is positive. For example, if *D* is 2×2 with a negative trace, positive determinant, and one positive diagonal entry and the equilibrium point is a CVS, then the equilibrium is an evolutionary branching point. Alternatively, if all diagonal entries of *D* are negative and the CVS condition is satisfied, then the equilibrium point is an attractor and evolutionary branching does not occur.

**Table A3:** Relations between subsystem stability, CVS, and evolutionary branching conditions for frequency independent selection.

|  | CVS condition |  |
| --- | --- | --- |
| | $s(D + CA^{-1}(-B)) < 0$ | $s(D + CA^{-1}(-B)) > 0$ |
| Single trait |  |  |
| $d > 0$ | branching point | repellor |
| $d < 0$ | CVS | repellor |
| Multiple traits |  |  |
| $D_{ii} > 0$ for some $i$ | branching point | repellor |
| $D_{ii} < 0$ for all $i$ | CVS | repellor |

### A4. DETAILS OF MODEL AND NUMERICAL ANALYSES FOR TWO COMPETING SPECIES

In this appendix, we provide the details on the models and numerical analyses for the figures, which were done in R (R core development team 2013) using packages deSolve (Soetaert et al. 2010) and rootSolve (Soetaert 2009).

For the stability bifurcation analysis in Figure 5 in the main text, i.e. competition model I, we reproduced the exact model from (Vasseur et al. 2011). We copy their model into this appendix to make it easier to reference. Vasseur et al. (2011) model the dynamics of the densities of two competing species, *N*_1_ and *N*_2_, and the mean trait of species 1, *x* as

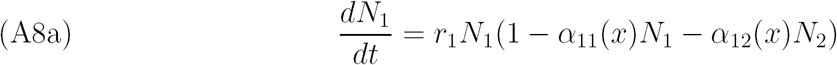

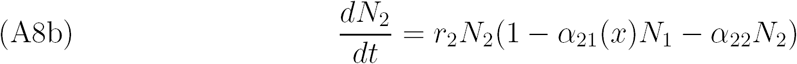

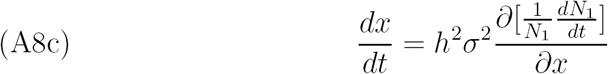

where the interspecific competition between species and intraspecific competition within species 1 depends on the mean trait *x* as follows

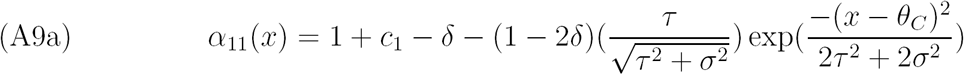

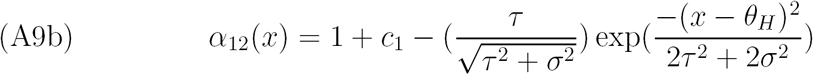

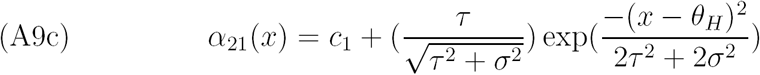

These curves are plotted in Figure 4 I in the main text with parameters *θ*_*C*_ = 0, *θ*_*H*_ = 1.6, *τ* = 0.5, *σ* = 0.1, *c*_1_ = 0.1, *δ* = 0.4.

In Figure 5, parameters are as follows: *r*_1_ = 3, *r*_2_ = 1, *τ* = 0.5, *σ* = 0.32, *c*_1_ = 0.1, *δ* = 0.4, *θ*_*C*_ = 0 and *θ*_*H*_ varied from 0.75 to 1.75, while *α*_22_ varied from 0.1 to 0.9, both in increments of 0.001. Varying *θ*_*H*_ (trade off) alters the evolutionary stability, while varying *α*_22_ (intraspecific competition of species 2) alters the ecological stability. Note that the heritability (*h*) in (A8) separates the time scale of the evolutionary process from the ecological and importantly, has no effect on the equilibrium values. To determine stability for slow and fast evolution for each point, we found the equilibrium point and evaluated our stability conditions. Then, for the regions with transitions in stability, we varied the heritability and evaluated stability from the full Jacobian. In Figure 5A, the gradient shows the critical heritability at which the transition in stability occurs. For plots *B* and *C*, we numerically solved the model while varying heritability from 0 to 1 with *α*_22_ = 0.75, 0.5 and *θ*_*H*_ = 1.5, 1.0, respectively, and found the minimum and maximum population densities from 9000 to 10000 time steps.

For competition model II, we slightly modified the model from Vasseur et al. 2011. In particular, we modified *α*_21_ so that antagonistic effects of species 1 on 2 increase with the antagonistic effects of species 2 on 1 as follows

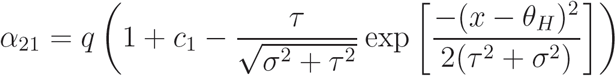

with introduced parameter *q* that represents how the antagonistic effects from species 1 on 2 scale with the effects of 2 on 1.

By varying the same parameters as in the test for traits in model I over a similar range, we tested for ecological, evolutionary and overall stability (Figure A1). Specifically, parameters are as follows: *r*_1_ = 3, *r*_2_ = 1, *τ* = 0.5, *σ* = 0.6, *c*_1_ = 0.1, *δ* = 0.4, *q* = 0.8, *θ*_*C*_ = 0 and *θ*_*H*_ varied from 0.5 to 1.9, while *α*_22_ varied from 0.5 to 0.9, both in increments of 0.01. In this range, there were always two equilibria, one of which that was always ecologically, evolutionarily, and overall stable. The stability properties of the other equilibrium depended on the parameter values, which partitioned equilibria into four categories, as in the analysis of trait model 1, with greater intraspecific competition in the non-evolving species leading to ecological stability and greater trade offs leading to evolutionary instability. This equilibrium was always overall unstable (Figure A1). In this bifurcation analysis, we needed to smooth the plots due to numerical difficulty in finding equilibrium points as the equilibrium approached the transition between ecological stability to instability and this led to the jagged curves.

**FIGURE A1.**
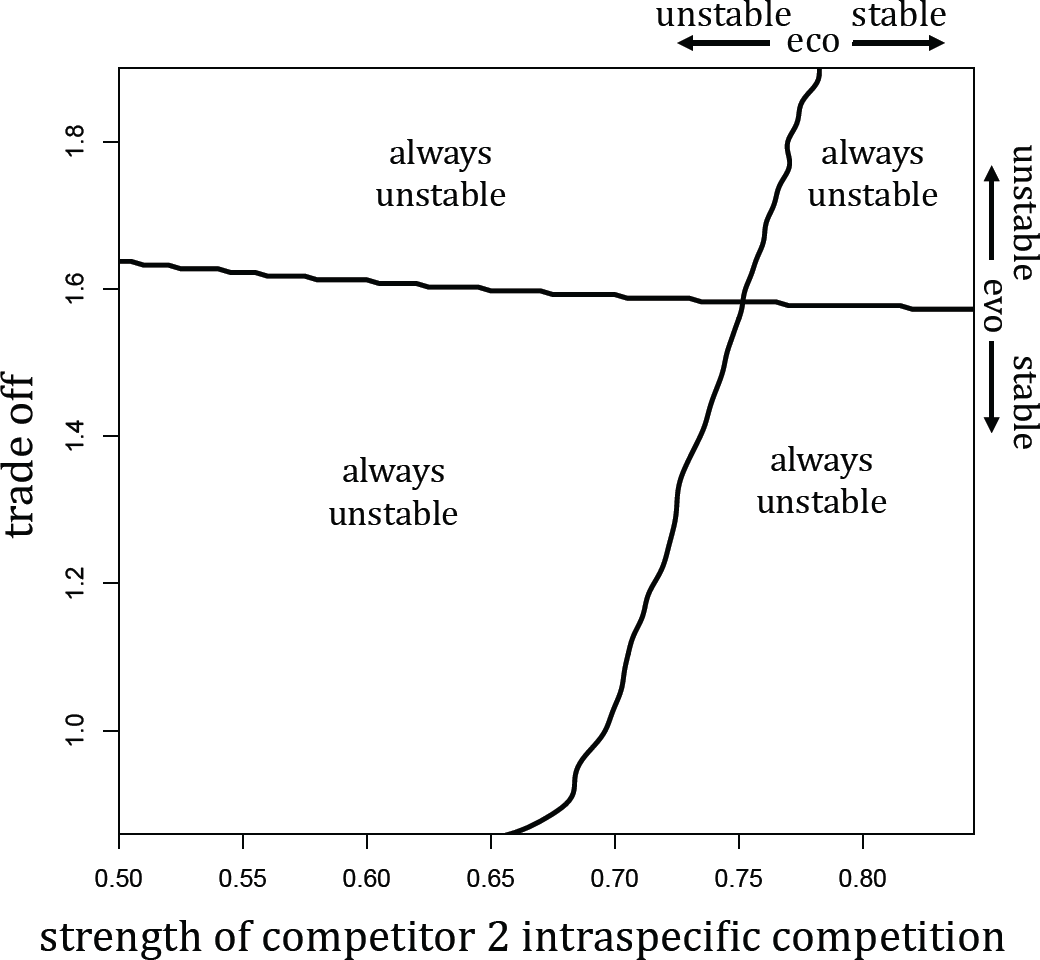
Bifurcation plot showing the ecological stability, evolutionary stability and overall stability for various relative rates of evolution for Model II. There were always two equilibria in this region. One was always ecologically, evolutionarily, and eco-evolutionarily stable. Here, we only show the other one, which was always eco-evolutionarily unstable.

### A5. DETAILS OF MODEL AND NUMERICAL ANALYSES FOR FOOD CHAIN

In this appendix, we describe in more detail the food chain model and the numerical methods we use to determine the effects of eco-evolutionary feedbacks. In particular, we explain how we generated the figures in this section.

Our three-species food chain is composed of a basal resource with density *N*_*y*_, an herbivore with density *N*_*x*_ and a top predator with density *N*_*z*_. We model the herbivore evolving in two correlated traits, with means *x*_*i*_, *i* = 1, 2. Trait one and two determine the interactions of the herbivore with the resource and the predator, respectively. Higher values of trait one increase the attack rate on the herbivore, while higher values of trait two increase defense against the predator. Both traits trade off with intraspecific competition. Finally, we assume that the traits independently contribute to herbivore fitness. It is important to note that because such a system is degenerate for perfectly positively or negatively correlated traits, our theory is only valid for a range of correlations within (*−*1, 1).

We use this information to determine the sign structure for the eco-evolutionary feedbacks for slow and fast evolution. First, our food chain model has a characteristic sign structure for the ecological matrix

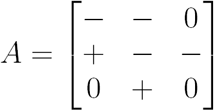

From classic Lotka-Volterra food chain theory, when an equilibrium exists, it is always ecologically stable (*s*(*A*) *<* 0, Hofbauer and Sigmund (1998)). Recall that from (3) in the main text, if genetic variances of both traits are equal and we let *ϵ* = *v*^2^, we can express 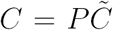, where *P* is the correlation matrix. Under the assumptions of the two traits, the signs of matrices *B* and 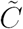 are

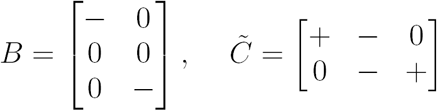

Assuming an equilibrium is ecologically stable, altogether, this leads to the eco-evolutionary feedback via the evo-eco-evo pathway

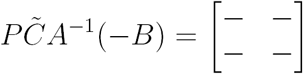

Adding this matrix to matrix 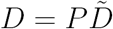 will make the trace more negative, but have undetermined effects on the determinant. Hence, the feedback can have stabilizing or destabilizing effects on the equilibrium.

In the following analysis, for when evolution is fast, we assume that the equilibrium is evolutionarily stable (since otherwise, the equilibrium would be overall unstable independent of the eco-evolutionary feedback). Under this assumption, 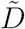 has sign structure

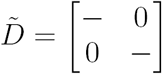

From this we can infer that the feedback via the eco-evo-eco pathway is

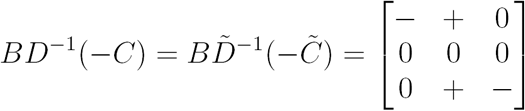

Note that *P* disappears from the expression of this feedback. Adding this to matrix *A* will make the trace more negative but again have undetermined effects on the determinant. Hence, overall, the effects of eco-evolutionary feedbacks on stability cannot be determined purely on sign structure.

To explicitly test for the effects, we used packages rootSolve (Soetaert 2009) and deSolve (Soetaert et al. 2010) in R to find equilibria, evaluate the Jacobian, and numerically solve, the following model

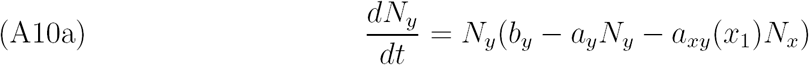

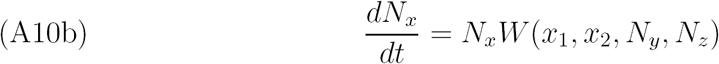

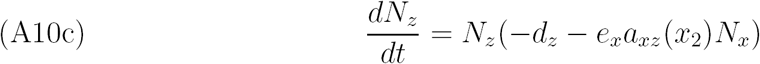

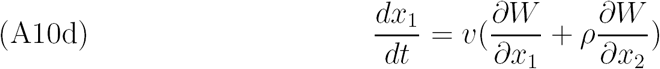

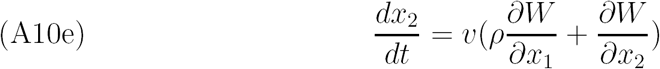

where *b*_*y*_, *a*_*y*_, *d*_*z*_ and *e*_*x*_ are the resources intrinsic growth rate, the resources intraspecific competition, the top predators intrinsic death rate, and the efficiency with which the top predator converts the herbivore into its own offspring, respectively. In the evolutionary dynamics, *v* and *ρ* are the genetic variance of both traits and the correlation, respectively. Note that equilibrium values are independent of *v* and *ρ*. In the model,

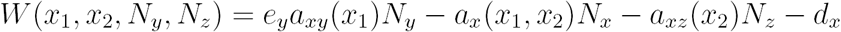

is the fitness of the herbivore as a function of mean traits *x*_1_ and *x*_2_, with *e*_*y*_ as the efficiency with which the herbivore converts the resource into offspring and *d*_*x*_ is the herbivores intrinsic death rate. Here,

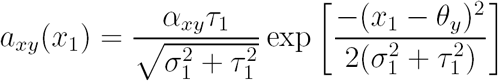

is the mean attack of the herbivore population on the basal population as a function of mean trait *x*_1_ (Schreiber et al. 2011), and

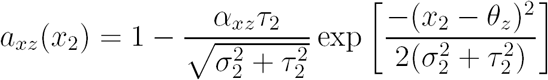

is the mean attack of the predator population on the herbivore population as a function of defense trait *x*_2_ and

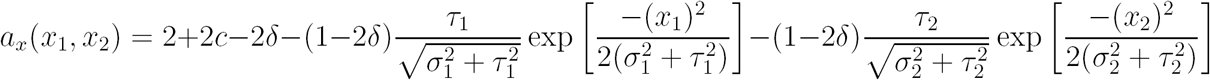

is the mean total intraspecific competition as a function of mean traits *x*_1_ and *x*_2_. To arrive at this expression, we used the same functional form from Vasseur et al. (2011) that describes how intraspecific competition depends on each trait and then summed these together. The additional parameters are described in Vasseur et al. (2011).

For Figure 6, we used the following parameters 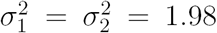, *τ*_1_ = *τ*_2_ = 1.5, *θ*_*y*_ = 4.1, *θ*_*z*_ = 5.5, *α*_*xy*_ = *α*_*xz*_ = 0.8, *a*_*y*_ = 0.3, *b*_*y*_ = 0.65, *d*_*x*_ = 0.5, *d*_*z*_ = 0.1, *e*_*y*_ = *e*_*x*_ = 0.9, *c* = 0.1, and *δ* = 0.01. In (A), we varied *ρ* from *−*0.9999 to 0.9999 and *v* from 0.099 to 9.9 and calculated the stability modulus of the Jacobian. In (C), we plotted the minimum and maximum population densities and traits between time 2000 and 3000 from the numerical solutions of (A10) for *v* = 1.98 for *ρ ∈* (*−.*9,.9).

